# SARS-CoV-2 genome analysis of strains in Pakistan reveals GH, S and L clade strains at the start of the pandemic

**DOI:** 10.1101/2020.08.04.234153

**Authors:** Najia Karim Ghanchi, Kiran Iqbal Masood, Asghar Nasir, Waqasuddin Khan, Syed Hani Abidi, Saba Shahid, Syed Faisal Mahmood, Akbar Kanji, Safina Razzak, Zeeshan Ansar, Nazneen Islam, M. B. Dharejo, Zahra Hasan, Rumina Hasan

## Abstract

**Objectives:** Pakistan has a high infectious disease burden with about 265,000 reported cases of COVID-19. We investigated the genomic diversity of SARS-CoV-2 strains and present the first data on viruses circulating in the country.

**Methods:** We performed whole-genome sequencing and data analysis of SARS-CoV-2 eleven strains isolated in March and May.

**Results:** Strains from travelers clustered with those from China, Saudi Arabia, India, USA and Australia. Five of eight SARS-CoV-2 strains were GH clade with Spike glycoprotein D614G, Ns3 gene Q57H, and RNA dependent RNA polymerase (RdRp) P4715L mutations. Two were S (ORF8 L84S and N S202N) and three were L clade and one was an I clade strain. One GH and one L strain each displayed Orf1ab L3606F indicating further evolutionary transitions.

**Conclusions:** This data reveals SARS-CoV-2 strains of L, G, S and I have been circulating in Pakistan from March, at the start of the pandemic. It indicates viral diversity regarding infection in this populous region. Continuing molecular genomic surveillance of SARS-CoV-2 in the context of disease severity will be important to understand virus transmission patterns and host related determinants of COVID-19 in Pakistan.

## Background and Rationale

The global outbreak of the novel coronavirus 2019, SARS-CoV-2 (causative agent of COVID-19), has caused over 566,654 deaths as of 13 July 2020, with more than 7 million individuals infected globally. SARS-CoV-2 was first reported in Wuhan, China associated with acute respiratory infection (1). The SARS-CoV-2 RNA virus genome is about 28.9 kb in size and phylogenetic analysis shows it to belong to the *Sarbecovirus* subgenus of *Betacoronavirus* and the family *Coronavirdae* (2). Its sequence is closely related to the Bat CoV RaTG13 with a sequence similarity of 96.3% across the genome. The virus has a higher rate of infectivity than previous coronaviruses such as the MERS (Middle East Respiratory Syndrome) virus and SARS (Severe Acute Respiratory Syndrome) virus outbreaks in 2014 and 2006 respectively (3).

SARS-CoV-2 is transmitted via droplets and small particles produced during coughing or sneezing and affects the upper respiratory tract including nose, throat, pharynx and lower respiratory tract (4). Most individuals present with fever, body ache, non-productive cough and shortness of breath, with a small proportion developing more severe disease including death. The case fatality rate (CFR) of SARS-CoV-2 infections has varied regionally. In China, the CFR ranged from 5.8 % in Wuhan to 0.7 % in the rest of the country. CFR at the peak of the pandemic in Mar was variable at 6.2% in Italy, 3.6% in Iran and 0.79% in South Korea (5). In later months, April to June, the CFR continued to vary globally. Reliable assessment of CFR may be limited by the degree of testing in the population however, the possibility of SARS-CoV-2 diversity may play a role in disease variability.

Since first identified as the Wuhan-1 strain in Hubei, China in January 2020, (6) SARS-CoV-2 strains have acquired mutations during its spread across regions that differentiate it into different global clades (7) (8). Initial SARS-CoV-2 strains were primarily L clade which subdivided to S and also sub-divided to V and G, as found in Asia, Oceania, Europe, South America, and North America; with clade I identified more recently. The GISAID database has made available greater than 63,000 SARS-CoV-2 global isolates (http://www.gisaid.org) to facilitate a comparison of strains.

Pakistan has a population of about 200 million with limited health resources. Karachi is its most populous city with about 20 million individuals. Up until mid-July approximately 262,000 COVID-19 cases have been diagnosed of which about 70,000 were from Karachi. Reduced rates of COVID-19 have been attributed to limited testing for SARS-CoV-2. Thus far, the case fatality ratio for SARS-CoV-2 in Pakistan has been 2% with some regional variations (9). Data from Pakistan regarding viral transmission has not been available. We sequenced SARS-CoV-2 isolates from Karachi and compared from Pakistan, global and Asian strains to understand transmission dynamics in the country.

## Materials and Methods

### Ethical approval

This study was approved by the Ethical Review Committee at the Aga Khan University (AKU), Karachi, Pakistan.

### Diagnostic testing for SARS-CoV-2

Nasopharyngeal swab specimens were tested for SARS-CoV-2 by reverse transcription (RT) polymerase chain reaction (PCR) at the Section of Molecular Pathology, Clinical Laboratory, AKUH. Specimens received in March were tested using the WHO protocol for the 2019-nCoV RT-PCR assay (10). Specimens in May were tested using the Cobas^®^ SARS-CoV-2 RT-PCR assay (Roche Diagnostics, USA). Respiratory samples archived at the Clinical Laboratory, Section of Molecular Pathology were used for virus extraction and testing. Laboratory data was (including age, gender) was utilized where available.

RNA extraction, Library preparation, NGS sequencing, and Consensus Sequence generation RNA was extracted from eight respiratory samples positive for SARS-CoV-2 using the QIAmp Viral RNA Mini kit (Qiagen). Sequencing was performed as described previously (11). First-strand cDNA synthesis was performed with SuperScript III Reverse Transcriptase (SSIII), Thermo Fisher Scientific, USA. Briefly, 8 μl of viral RNA was mixed with 1 μl of 50 ng/μl of random hexamer and 1 μl of 10mM dNTPs. Then denatured at 65°C for 5 minutes. Then, 10X Buffer, 0.1 mM DTT, RNase Inhibitor (40 U/μl), 25mM MgCl_2_ and 200 U SuperScript III Reverse Transcriptase was added. Second strand cDNA was synthesized with DNA polymerase I, Large Fragment, Klenow (Invitrogen, USA).

The library preparation was performed according to the Nextera XT DNA Library Preparation kit (Illumina) using 1 ng DNA. Bead-based normalization of the libraries was carried out as recommended by the manufacturer. Normalized libraries were equimolarly pooled and spiked with PhiX control prior to sequencing.

Sequencing was performed on the Illumina Miniseq platform using a 300 cycle Miniseq Reagent Kit v2 (Illumina). DRAGEN RNA Pathogen Detection App v3.5.7 on BaseSpace (12) was used to filter out human sequence reads by combining human (hg38) + SARS-CoV-2 virus reference (NC_045512.2) genome to generate consensus sequences (Supplementary Table 1). Quality metrics of genome data generated are provided in Supplementary Table 2.

### Variant calling and Phylogenetic analysis

FASTQ files were aligned to the SARS-CoV-2 virus reference genome Wuhan-1 (NC_045512.2) by BWA (13). PICARD tools (http://broadinstitute.github.io/picard/) were used to remove redundant alignments and calculating alignment statistics. Variants were identified by Genome Analysis Toolkit (GATK) (14). The effect on protein-coding by the mutation is determined by an impact score (15).

For phylogenetic analysis, 7 SARS-CoV-2 sequences from our study, 3 Pakistani SARS-CoV-2 sequences (Supplementary Table 1) along with the 449 full-length SARS-CoV-2 reference sequences (Supplementary Table 3) from different pandemic countries obtained from the NCBI SARS-CoV-2 Resources (https://www.ncbi.nlm.nih.gov/sars-cov-2/) were subjected to Multiple Sequence Alignment (MSA) along using MAFTT online server (16). The MSA was subsequently used to generate a Maximum Likelihood (ML) phylogenetic tree using PhyML 3.0 (http://www.atgc-montpellier.fr/phyml/) with a GTR-based nucleotide substitution model and aLRT SH-Like branch support. The root of the tree and branch length variance was determined using the TreeRate tool (17) by applying a generalized midpoint rooting strategy. The tree was visualized and edited in Figtree software (http://tree.bio.ed.ac.uk/software/figtree/). The mean and individual pairwise distance between 7 SARS-CoV-2 sequences from our study and 3 previously deposited Pakistani SARS-CoV-2 sequences was calculated using MEGA 7 (18).

For genomic epidemiology of Asian-strains focused subgroup analysis of SAR-CoV-2 as of 16^th^ July 2020, we downloaded 5,215 complete sequences of SARS-CoV-2 along with the required metadata from the GISAID considering the following parameters: 1) genome length > 29.0 bps, 2) further assigns labels of high-coverage <1% Ns – undefined bases, and 3) low-coverage >5% Ns. After the inclusion of our 7 strains to the fasta file, phylogenetic tree reconstruction was performed using NEXTSTRAIN’s (https://www.nextstrain.org/) augur (https://www.docs.nextstrain.org/projects/augur/en/stable/) pipeline. Ancestral state reconstruction and branch length timing were performed with TreeTime (19). Finally, the collection of all annotated nodes and metadata was exported to the interactive phylodynamic visualizing tool Auspice’s (https://www.nextstrain.github.io/auspice/) JSON format.

## Results

We obtained seven full genome sequences of SARS-CoV-2 strains and one partial sequence (Supplementary Table 2). All eight isolates were from individuals with mild COVID-19. Five strains sequenced isolates were travelers from Iran and Turkey in March, at the start of the COVID-19 pandemic in Pakistan. Three strains isolated from cases in May were from a traveler to Iran, a case of local transmission and a religious pilgrim who attended regional event. We further compared three Pakistani isolates present in the NCBI database which were from a traveler to Iran, a case of local transmission and one in which travel history was not available. Clinical information was not available for these cases. Overall we had data for 10 strains; from seven male and three female cases. The age group of infected individuals based on available data was n=2, less than 18 y; n=4, 19-35 y; n=2, 36-50 y and n=2, > 50 y.

### Phylogenetic analysis

Initial phylogenetic analysis was done using ten SARS-CoV-2 strains from Pakistan (Supplementary Table 2) as compared with 449 global isolates from different pandemic countries obtained from the NCBI SARS-CoV-2 Resources (https://www.ncbi.nlm.nih.gov/sars-cov-2/) (Figure 1). Two strains (S2 and S3) clustered with those from Saudi Arabia, and India (Figure 1). S5 clustered with a strain from the USA. S19 clustered with a strain from Australia. S10 and S11 clustered with strains from Saudi Arabia. S21 was related to strains clustered from the USA and Australian. Additionally, Pakistani strain analysis revealed PAK/KHI1 and PAK/Gilgit1sequences matched with Chinese isolate and PAK/Manga1 with an isolate from the USA.

**Figure 1.**
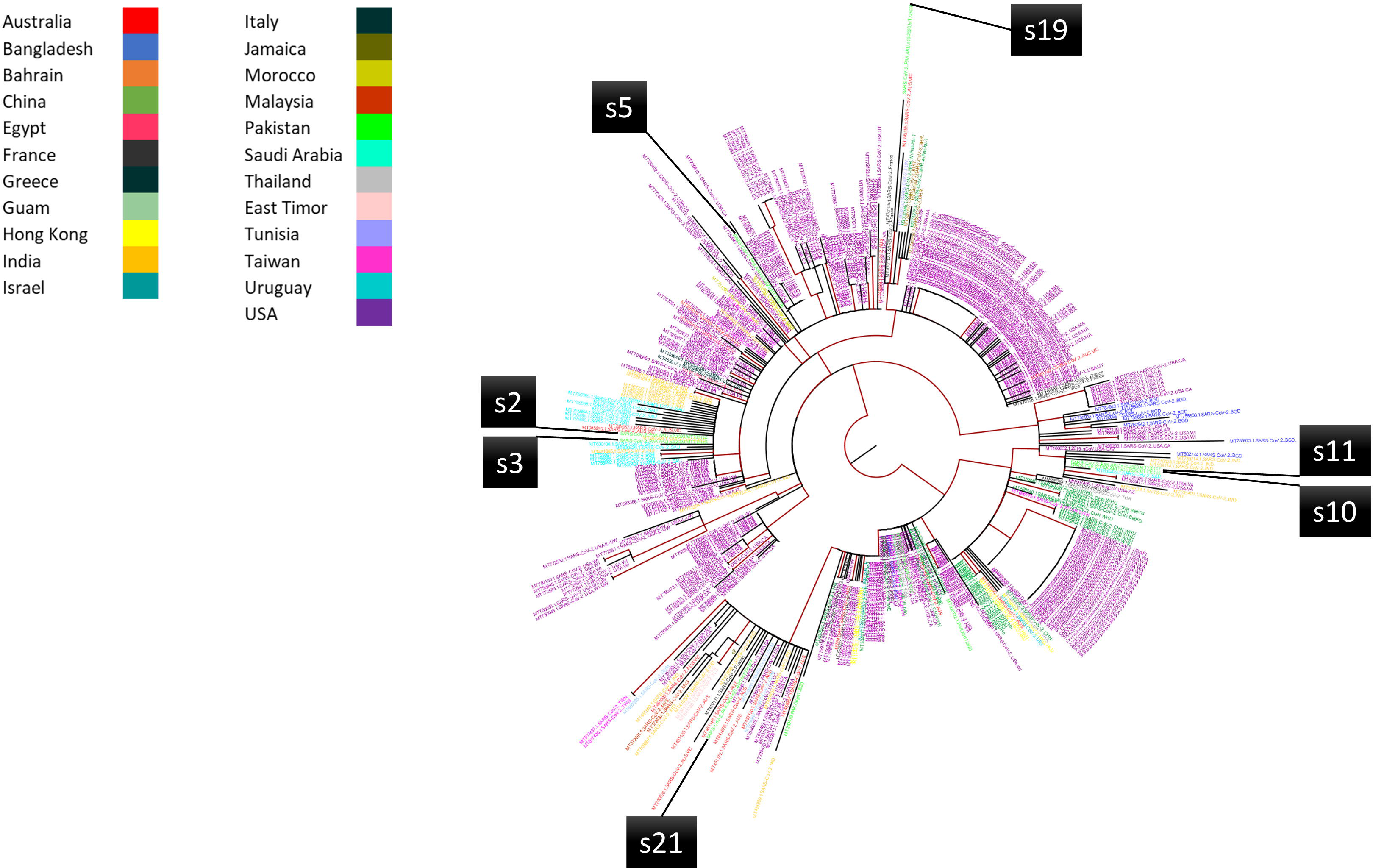
Maximum-Likelihood phylogenetic tree of SARS-CoV-2 sequences from Karachi. The tree was constructed using Karachi sequences (Supplementary Table 2) along with the 449 full-length SARS-CoV-2 reference sequences (Supplementary Table 3). Karachi sequences are shown in green colour. The root of the tree was determined using TreeRate tool by applying generalized midpoint rooting strategy. The tree was visualized and edited in Figtree software.

The mean pairwise genetic distance between our sequences; sequences previously deposited from Pakistan, and sequences from India, Saudi Arabia, USA, and Australia was found to be 0.00, indicating phylogenetic relatedness between these sequences.

We further expanded the genomic epidemiological analysis by focusing on 5,215 Asian isolates available in the GISAID database. As depicted in Figure 2, in the absence of non¬Asian sequences, sequences S2 and S3 clustered with the Japanese sequences (Supplementary Table 1). S5 clustered with a strain from Israel; S10 and S11 clustered with strains from Saudi Arabia. S19 matched a Pakistani isolate with the divergence rate of 0.0007953 and was a case of local transmission. S21 was most closely related to a strain from India. Additional Pakistani strain analysis revealed; PAK/KHI1 matched an Indonesian isolate, PAK/Gilgit1 with a Chinese isolate and PAK/Manga1 with an isolate from Singapore.

**Figure 2.**
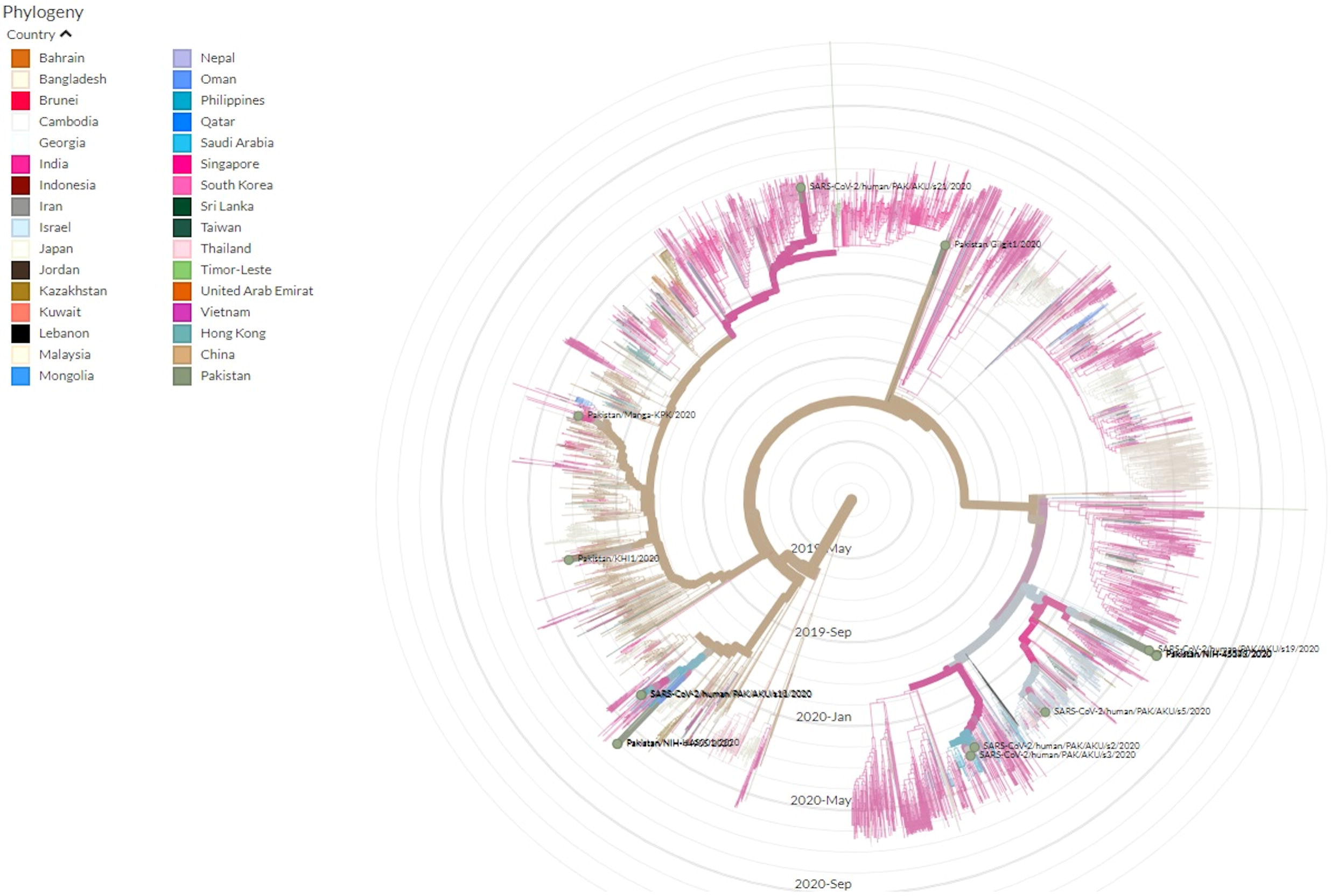
Time-resolved phylogenetic distribution of genomic epidemiology of SAR-CoV-2 focused on Asian subsampling. (Screenshot of the current Nextstrain display in SVG format are protected by CC-BY license). Tree option layout is selected as radial, branch length is set as time interval while branch labels as amino acid substitution **(A)** Asian subsampling, and **(B)** Asian subsampling with highlighted Pakistani SARS-CoV-2 isolates (total 16, 7 are from this study inclusive).

We further analyzed genome sequences of these ten full and one partial (S16) SARS-CoV-2 isolate. Of the eleven, five were G clade of the sub-clade GH; three were from March (S2, S3, and S5 from Iran and Turkey travelers) and two from May (S16 from an Iran traveler and S19 through a local transmission). Two isolates from travelers to Turkey (S10) and Iran (S11) in March belonged to the S clade. Three strains from May comprised two GH clade strains, one from an Iran traveler (S16) and the other from local transmission belonged to L clade, with 99% homology to the Wuhan-Hu-1 nCoV strain type. Of the three Pakistani isolates present in the NCBI database, PAK/KHI1 belonged to L clade, PAK/Gilgit-1 from a traveler to Iran belonged to I clade and PAK/Manga1 was an L clade strain transmitted locally.

### Mutation Analysis

Variant analysis of the genomes revealed 34 SNV comprising of 2 non-coding, 18 non-synonymous and 14 synonymous variants (Table 1). The Orf1ab gene region exhibited the highest number of variants in all isolates, with 19 different variants (Table 2). Two Orf1ab mutations associated with evolutionary changes at nucleotide positions 8782 (2839S) and 14408 (P4715L) were observed. Variation in Orf1ab-8782S and Orf8-L84S are associated with evolutionary changes dividing SARS-Co-V-2 into lineages L and S (20) of which, the L lineage is more prevalent. We found S clade strains (S10, S11) with ORF8 L84S and ORF1ab 2839S mutations. These two strains also had S202N in the nucleocapsid and 302T in Spike gene regions.

**Table 1.**
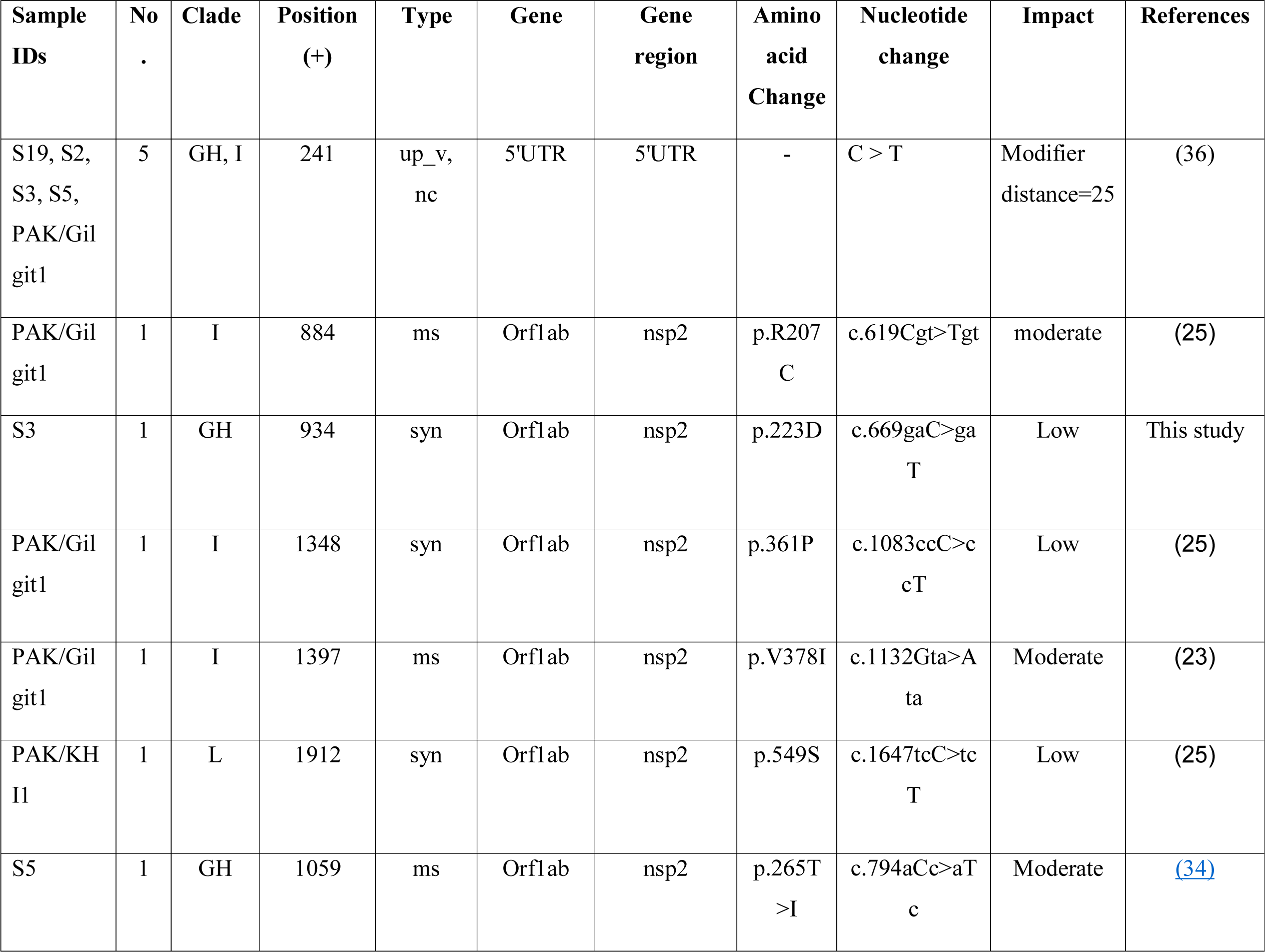
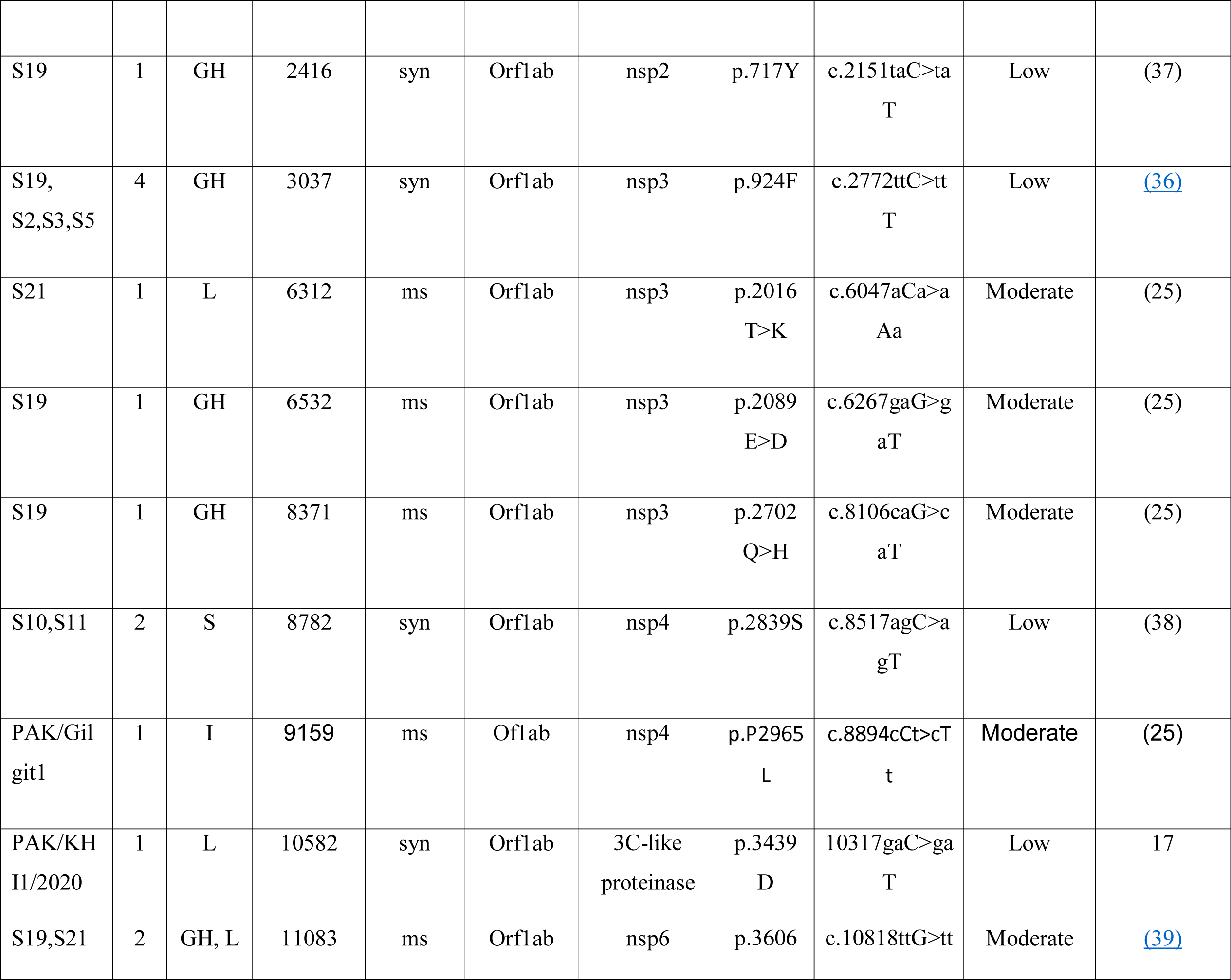
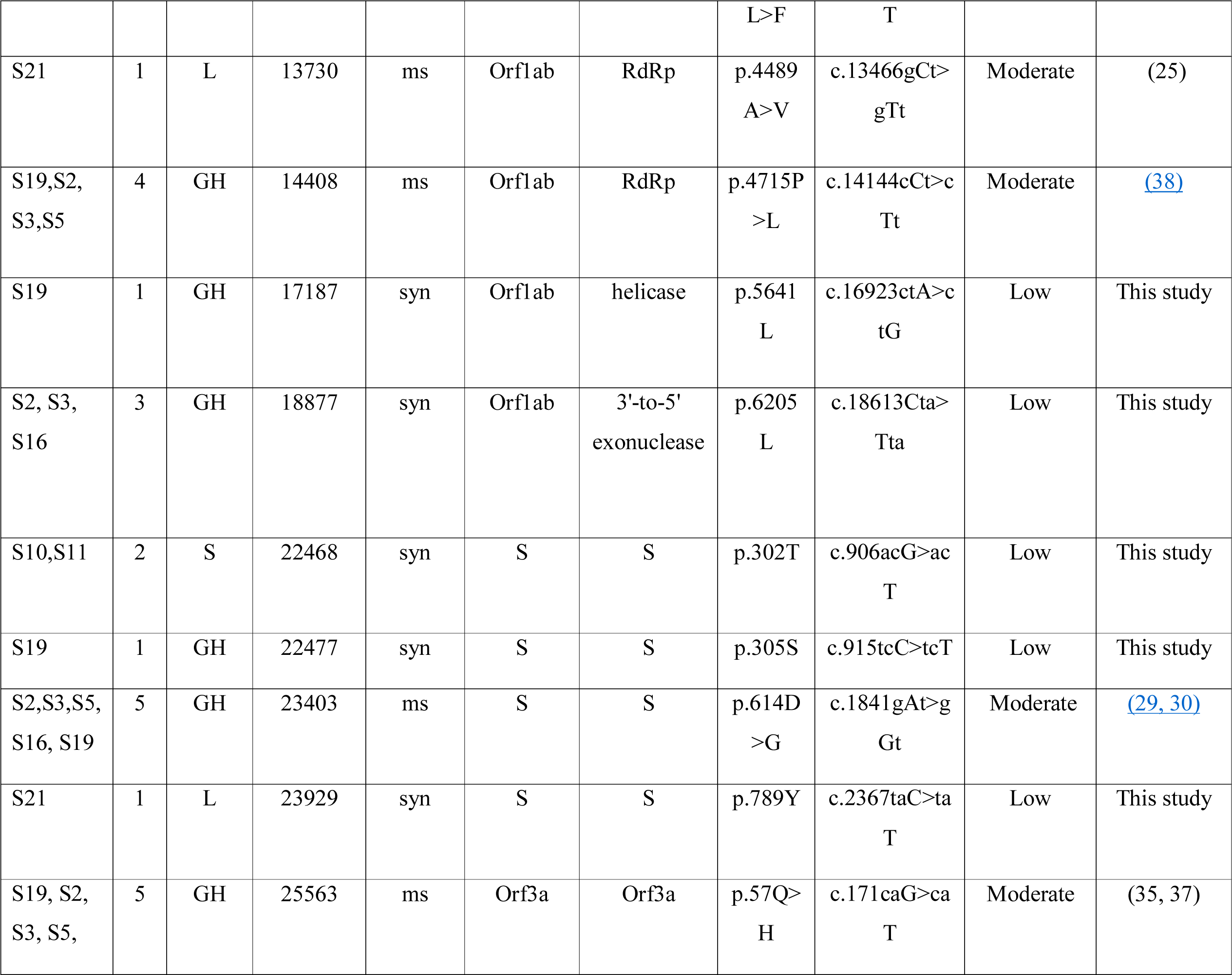
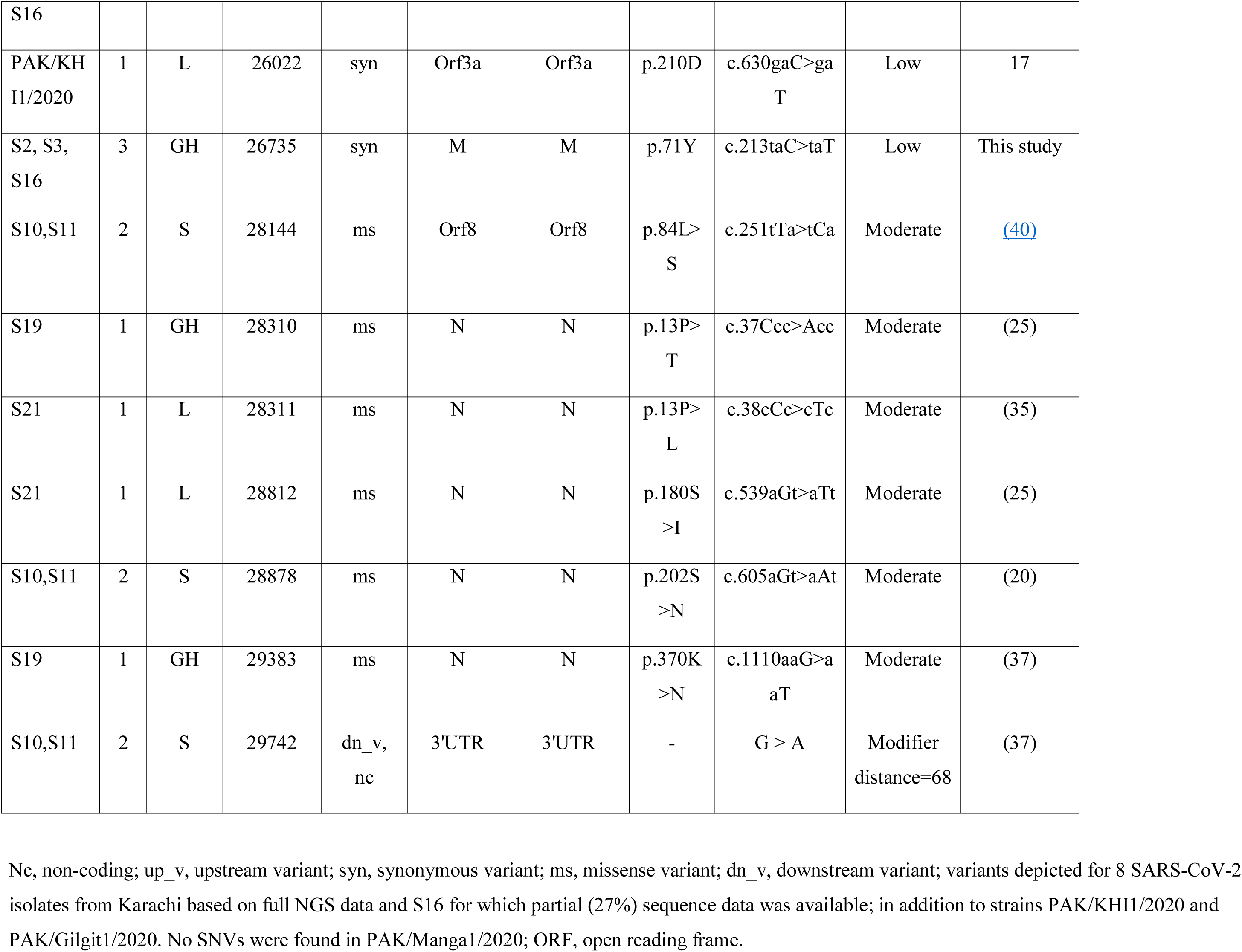
Description of single nucleotide variants (SNV) found in SARS-CoV-2 isolates from Karachi

**Table 2:**
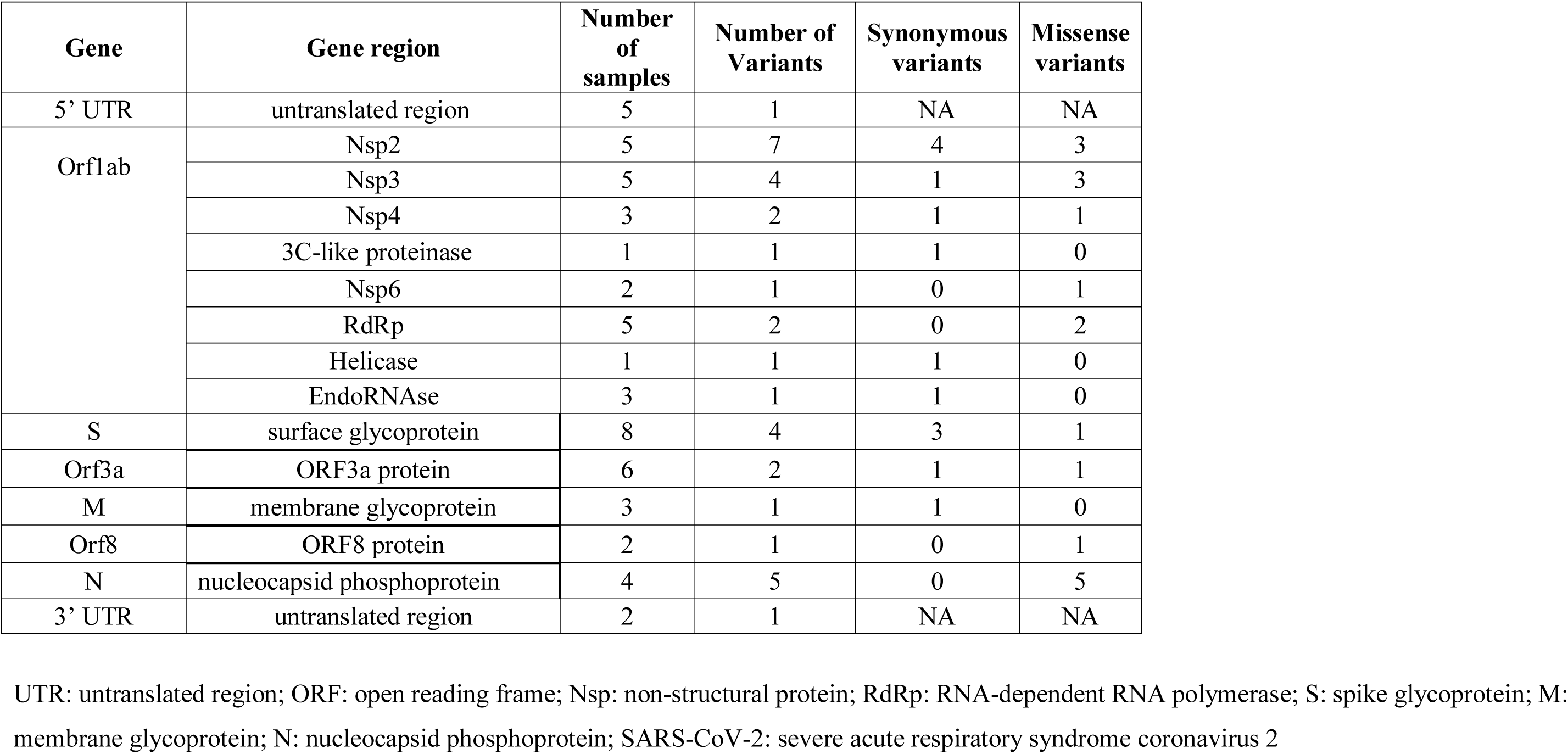
Number of variants detected in the gene regions of SARS-CoV-2 virus

The Spike glycoprotein mutation D614G defines the virus clade ‘G’ which further splits into GH and GR clades (20). D614G with ORF3a-Q57H found in five isolates (S2, S3, S5, S16, S19) identified these as GH clade. In four of the GH isolates we found P4715L/P323L in the nsp12 protein. We found GH clade strains to have additional Orf1ab variants at positions T265I, E2089D and Q2702H. One GH (S19) and one L (S21) clade strain both had the Orf1ab variant L3606F, which is associated with divergence of L clade towards the S clade and also has been shown as an additional mutation found in G clade strains (7). The L clade (S21) isolate had an additional Orf1ab mutation in the nsp4 region producing the A4489V change.

Of the three SARS-CoV-2 isolates previously submitted in the NCBI database from Pakistan, one (PAK/Gilgit1) had ORF1ab-encoded non-structural protein 2 region mutation G1397A producing a non-synonymous change V378I mutation. This mutation has been used to identify strains belonging to clade IV or I lineage isolates (21, 22). Further, this isolate had four other variants in Orf1ab; two were synonymous and two were non-synonymous, R207C and P2965L. The other two Pakistani isolates present in the NCBI database belonged to the L clade closely resembling the Wuhan-Hu-1 strain. One strain had three synonymous variants only and the other had genomic variants in the coding region.

Variants classified as having a moderate effect on the SARS-CoV-2 protein structure were found in the nucleocapsid N region (Table 1). Five mutations were found in the N gene region in isolates S10, S11, S19 and S21 from S, GH and L clades, respectively. One GH clade strain had P13T and K370N, one L clade strain had P13L, S180I and we observed S202N in two S clade strains. This indicates variability in the N gene as shown previously across global isolates (23). Structural proteins E and M genes showed most conserved regions across all the isolates with a silent variant at position 71Y in M proteins found in three GH clade strains. No variants were observed in ORF10, ORF6, and ORF7a region.

## Discussion

This study provides insights into the SARS-CoV-2 strains circulating in Pakistan using a small data set with nine genomes from Karachi and two from Northern Pakistan. We demonstrate the presence of SARS-CoV-2 GH, S, I and L clade strains in March at the start of the epidemic. The presence of different clades is not surprising as initial strains were all associated with travelers who presented with symptoms. The import of cases from locations such as Iran, UK and Turkey in March while the pandemic was spreading globally provides an opportunity for variable strain types to be studies.

Considering S2 and S3 which were GH clade strains, global phylogenetic analysis paired these with strains from Saudia Arabia whilst Asian subsampling showed clustering with Japanese sequences, which might indicate that strains from these countries might be genetically similar and might be evolving at a similar rate.

S19 clustered with an Australian isolate in comparison with global strains. In the Asian strain-based Nextstrain, S19 matched a local Pakistani isolate which correlated with it being a case of local transmission. Phylogenetic analysis with global sequences showed S21 to be related with USA and Australian strains, while Asian-focused real-time strain evolution, as inferred by the time-resolved phylogenetic tree analysis, showed phylogenetic relationship with an Indian strain. This isolate was from a religious pilgrim who attended an event in Punjab province which was subsequently associated with a super spread of SARS-CoV-2 infections. The period when the sample was collected further supports virus evolution with the nearest strain is at the Indo-Pak region. S10 and S11 (S clade) exhibit coincident time lineages with Saudi Arabian strains; these were from travelers from Iran and Turkey who returned from religious pilgrimage and have acquired Saudi Arabian strains similar to the hCoV-19 Saudi Arabian strains.

Clade GH has been associated with returning travelers from Iran to other countries (22). We found GH clade strains to be associated with travelers from Iran and Turkey concurring with previous reports (22).

S21, a L lineage isolates, found to be closest to the Hubei strain, was from a religious pilgrim who had attended an event in Punjab Province where it is believed that SARS-CoV-2 infections were spread through attendees who had traveled from China. This may explain similarity of the isolate with the Wuhan-Hu-1 SARS-CoV-2 reference strain. This L lineage isolates exhibited G11083T mutation which was first reported on January 2020 in China and is associated with super spreader events in the USA, Singapore, Japan, and Europe (7). The L clade further evolved into the V clade with the emergence of G26144T. L clade is highly variable and shown to harbor up to 12 SNVs in the early phase of the COVID-19 pandemic. Further, GH and L clade strains in our cohort were also identified from both overseas travel and local transmission events.

Of the two SARS-CoV-2 evolutionary lineages L and S, the L lineage continued to split initially equally into G and V versions, with G reaching 50% of isolates in March 2020 and then splitting further into GR and GH subclades (24). G clade strains are identified by the Spike protein D614G mutation and then sub-divided into the H clade with the ORF3a Q57H mutation (25). We found that S19, a GH clade, and S21, an L clade strain both had Orf1ab L3606F mutation. S19 in addition to having GH clade mutations D614G, Q57H, and P323L, had an additional 4 mutations. Of note, both S19 and S21 were strains from May, later in our study set and the presence Orf1ab L3606F mutation suggests additional evolutionary changes in these isolates.

The spike glycoprotein facilitates SARS-CoV-2 entry into host cells by binding to the ACE2 receptor (26). The D614G mutation has been associated with increased virulence and transmission of SARS-CoV2 isolates, demonstrating a higher viral load within infected individuals (27). It is speculated that mutation to Glycine at the 614 site may introduce structural instability into the spike protein (28). *In vitro* studies using a mutant Spike protein with D614G have shown that the virus may be more infective due to the correlated reduction in viral shedding of the receptor binding S1 domain which is associated with the mutation, leading to increased S-protein incorporation into the virion (29). Thus far, G clade isolates have been found globally and are thought to comprise a high proportion of isolates in Europe and North America. It has been suggested that the S D614G mutation is associated with greater mortality observed in Belgium, Spain, Italy, France, Netherlands and Switzerland (30). However, it is difficult to draw conclusions from laboratory studies and the impact of the D614G variant on transmission between patients and across a population.

Variants in the non-coding 5’UTR and 3’ UTR regions of the SARS-CoV-2 virus have been reported to affect viral replication and transcription. TAR DNA binding protein (TARDBP) is a predominantly nuclear RNA/DNA-binding protein that functions in RNA transcription, splicing, transport, and stability. The variant (241C > T) has been reported to result in the strong binding of TARDBP to the 5’UTR region of the SARS-CoV-2 virus (31). This has been implicated in facilitating the translation of viral proteins resulting in its effective propagation within the human host. Interestingly, variant (241C > T) of the 5’UTR region often coexists with spike glycoprotein variant (S protein, D614G) (32). This coexistence is also evident in our study as we found 4 GH clade and an I clade strain to have the 5’ UTR variant +241 C>T.

Whilst the 2 S clade strains had the 3’ UTR + 29742 G>A. Human microRNAs (miRNAs) are noncoding RNAs that bind to complementary sequences in the 3’-UTR of the target RNAs and regulate the stability of the RNA at a post-transcriptional level. In addition, they also modulate different stages of viral replication, either positively or negatively. The variant (29742G>A) in the 3’UTR region of the SARS-CoV-2 virus has been reported to suggested to affect the binding of the miR-1307 and potentially causing a weakened host immune response against the virus (33). Hence, non-coding variants in both 5’ and 3’ UTR regions may enhance the virulence of SARS-CoV-2 strains however it is yet to be proven clinically.

The most common missense variants in the Pakistan SARS-CoV-2 strains were Orf1ab P4715L, Q57H and D614G which have previously been mainly reported from Europe and United States (23). Mutations were observed Orf1ab in all isolates studied and comprised thirteen different variants, exhibiting the highest mutability as shown previously (23). Two isolates (S10, Sll) had the S clade mutation ORF8 L84S and also the N gene S202N.

We found Orflab and N region to have the highest number of mutations as has been shown previously (34). The N protein mutation at Pro13 was present in both GH clade and L lineage isolates. This has mutation been reported as P13L previously in patients from the UK and Australia (35).

Overall, mutations in SARS-CoV-2 are an important mechanism to study variability and spread of the virus and should ideally be done in the context of the clinical disease caused by taking into account host factors as well. Our results are important as they highlight the diversity of SARS-CoV-2 strains at the start of the epidemic in Pakistan. The study was small due to resource constraints and the difficulty in access to NGS reagents in a global lock-down where shipments were stopped, it indicates that at the time of initial spread there were L, S, G and I clades circulating in the country. Initially, the epidemic was associated with travelers and local transmission of SARS-CoV-2 within a couple of weeks of the first known case. It would be of great importance to study the variability in the strains over the past months and also to see how they may differ between those associated with travelers and local transmission. We now hope to be a stage of flattening the COVID-19 curve with reducing rates of positive cases. It would be important to continue surveillance and conduct molecular epidemiological testing of the isolates to understand the disease in the context of strain diversity. This would also be important to understand local host immunity against the circulating SARS-CoV-2 and to investigate possible virulence characteristics associated with the changing viral genome and to relate these to disease severity in a country with a very high infectious disease burden.

## Acknowledgements

This study received funding support from Health Security Partners, USA and University Research Council, The Aga Khan University. We thank Drs. Roger Hewson and Barry Atkinson, Public Health England, UK for their assistance is establishing SARS-CoV-2 diagnostics. Thanks to the European Virus Archive - Global (EVAg), a European Union infrastructure project for making available control material for the study. Thanks for technical support to the Aga Khan University Hospital (AKUH) Clinical Laboratory sections of Molecular Pathology and Microbiology.

## Author contributions

Study design and funding (ZH, RH), Research methodology and Data collection (ZH, NG, KI, AN, SS, AK, ZA, NI, MBD), Data analysis (WK, SHA, NG, AN, ZH), Writing (ZH, RH, NG, AN, SFM)

